# *α*-tubulin acetylation and detyrosination correlate with starvation-induced autophagy in tobacco cells

**DOI:** 10.1101/111484

**Authors:** Dmytro I. Lytvyn, Alla I. Yemets, Yaroslav B. Blume

## Abstract

Recent data has enabled discovery of novel functions of microtubules (MTs) in the regulation of autophagy development under physiologic/stressful conditions in yeast and animal cells. MTs participate in maturation and traffic of autophagosomes through their dynamic state changes and post-translational modifications of tubulin, including acetylation. We demonstrated the involvement of tobacco cell MTs in the development of starvation-induced autophagy via tubulin acetylation and denitrotyrosination. Induced metabolic stress caused by prolonged cultivation of BY-2 suspension cells results in glucose depletion in the culture medium and following increased rates of protein hydrolysis and autophagy. Development of autophagy was strongly accompanied by α-tubulin acetylation and detyrosination. Both post-translational modifications were caused by changes in the molecular microenvironment of the tobacco cell MTs that was revealed via Co-IP assay. The termination of autophagy led to the development of programmed cell death that was characterised by nucleosomal DNA fragmentation and decreases in α-tubulin acetylation and detyrosination. Our data suggest the role of the functional state of MTs in the mediation of plant autophagy via changes in the tubulin microenvironment and in its post-translational modifications.

**SUMMARY STATEMENT:** The main findings cover a possible impact of plant microtubular cytoskeleton to starvation-induced autophagy development. It can be realized by means of tubulin post-translational modifications, acetylation in the first place.

## INTRODUCTION

Autophagy is known as a highly conserved degradation process present in all eukaryotes, including plants (Reumann et al., 2010; Avin-Wittenberg et al., 2012; Yoshimoto, 2012), that supplies nutritional recycling in the cell under developmental stages as well as under stress conditions (Liu and Bassham, 2012). The autophagic processes in plants are regulated by basically the same molecular machinery as in yeasts and animals because of their conservatism. It is provided by coordinated functioning of the ATG gene complex and can be separated into three groups: the ATG9 cycling system that controls lipid membrane recycling during autophagosome biogenesis; the PI 3-kinase complex that regulates an attraction of phosphatidylinositol 3-phosphate binding proteins to the preautophagosomal structure; and, finally, the ubiquitin-like Atg8 and Atg12 protein system that provides double-membrane enclosure through maturation of the autophagosome (Reumann et al., 2010; Liu and Bassham, 2012; Avin-Wittenberg et al., 2012). The paramount importance of this cellular process becomes apparent when cells are subjected to the impact of various external stressors such as oxidative stress, nutrition starvation, salt/drought stress and invasion by pathogens (Bassham et al., 2006; Liu et al., 2009; Han et al., 2011). In spite of the importance of autophagy for cell adaptation to adverse influence, however, many aspects of plant autophagy remain unclear.

On the other hand, to understand plant autophagy it is important to establish the role of other components of cellular machinery that mediate stress responses. In this regard, studies of the microtubular cytoskeleton deserve fixed attention when one considers the high sensitivity of this dynamic system as it reacts to external stimuli by regulating the functional state of the cell. It is known that plant microtubules (MTs) (as well as other eukaryotic MTs) are dynamically unstable structures formed by polymerisation/depolymerisation of α/β-tubulin dimers and involvement of numerous microtubule-associated proteins. This microenvironment determines cytoskeletal architecture and functional properties of MTs in general (Sedbrook, 2004; Hamada, 2007). The functions of MTs cover a wide range of cellular processes, including cellular growth, division and development as well as cell signalling and organelle transport (Wasteneys, 2004). On the other hand, the microtubular cytoskeleton is a highly receptive target for many external stimuli that substantially affect the dynamic structure of microtubules and thereby modulate cellular response. Heat shock, mechanical injuries, and cold stress (Hussey and Hawkins, 2001; Telewski, 2006; Sheremet et al., 2012; Nick, 2013) as well as salt stress (Wang et al., 2007, 2011) can be mentioned among the numerous abiotic factors that affect MTs. Moreover, the essential role of MTs in realisation of programmed cell death has also been shown in studies (Smertenko and Franklin-Tong, 2011). Other crucial reports put forth the novel view that MTs act as a signalling chain of autophagy development under both physiologic and stress conditions. MTs are important in key steps of autophagy including maturation and traffic of autophagosomes. Two functional characteristics of the MTs are crucial in these processes: their dynamic state and their tubulin post-translational modifications, including acetylation first and foremost (Monastyrska I., Rieter E., Klionsky et al., 2010; Mackeh et al., 2013). Some progress has been achieved in studies of MT functions in yeast and animal autophagy, but cellular events that implicate the MTs in autophagy development in plants remain to be clarified. This paper highlights the relationship and possible role of acetylation and detyrosination of α-tubulin in the development of starvation-induced autophagy in tobacco cells.

## RESULTS

### Glucose depletion in BY-2 culture and hydrolysis of cellular proteins

To characterize the metabolic stress taking place during prolonged cultivation, a comparative analysis was carried out of glucose depletion (as the main energy component of the growth medium) and changes in total protein concentration in cell suspension. Colorimetric determination of glucose concentration revealed its rapid lowering after day 5 of cultivation, which corresponded with the transition of the cell culture in the stationary growth phase. On day 7 of cultivation, the concentration of glucose was decreased to a critical value of 1.4 mg ml^-1^; the subsequent period (from days 8 to 11) was characterised by a complete lack of glucose in the culture medium by cultivation scheme 1 (Fig.1). At the same time, the dynamics of protein accumulation in the cell suspension appeared to be inversely proportional. The maximal protein concentration of 300 mg ml^-1^ occurred on day 5. During lengthier periods of time up to and including day 8, the protein concentration remained in the same range that corresponded to the stationary growth phase of the culture. From day 9 of cultivation forward, a gradual statistically significant decrease of protein concentration took place so that, during the last time period (day 11), the concentration was less than half the value of the maximum concentration (Fig. 1A). This dynamic decrease of protein concentration coincides in time with the sharp glucose depletion in culture medium and is probably the result of its exhaustion. A similar process was observed by application of another cultivation scheme 2 (diluted culture that grows up to 15 days inclusive). This experimental design of continuous cultivation was used to simulate a more prolonged starvation period to provide more pronounced cellular effects while reducing the amount of cell death. In this scenario, protein concentration was maximised from days 8-9 followed by protein hydrolysis (Fig. 1B) that was also caused by glucose depletion (not shown).

**Fig. 1.**
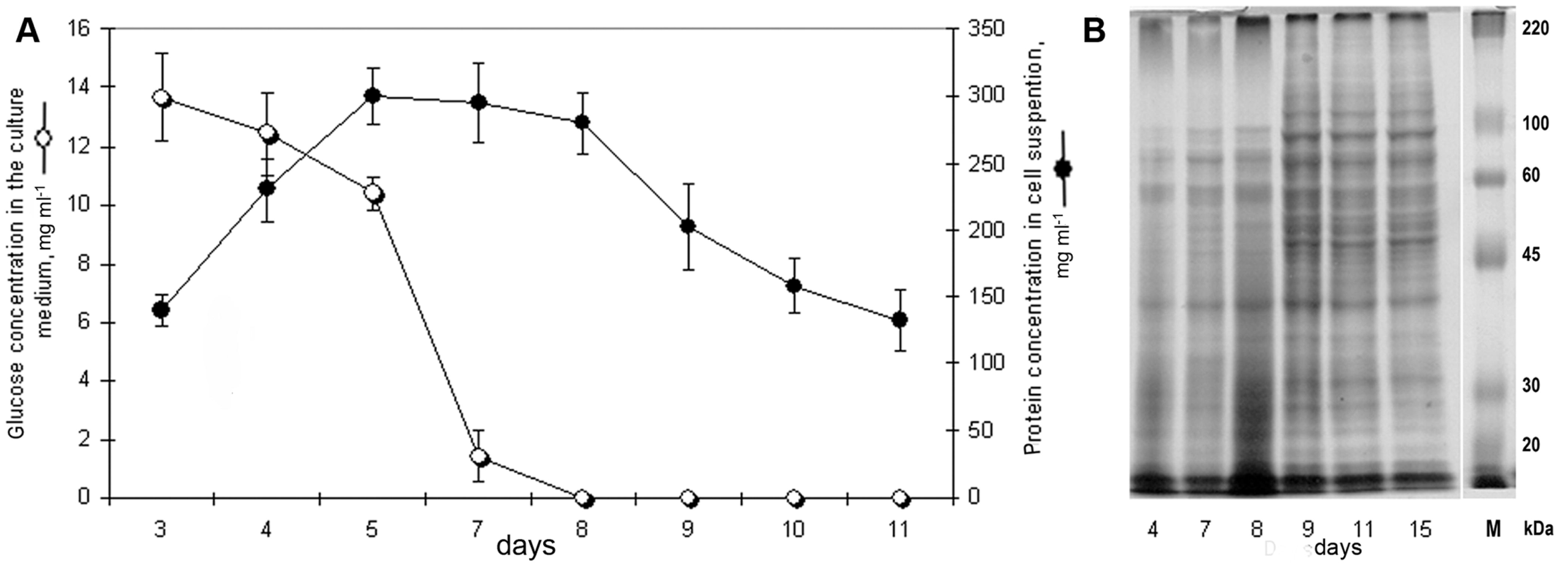
Impact of prolonged BY-2 cell cultivation on the induction of protein hydrolysis. (A) Correlations shown between glucose depletion and decreasing protein level. 11-day experiment (cultivation scheme 1). (B) SDS-PAGE of equal amount of BY-2 cell lysates from 15-day experiment (cultivation scheme 2), Coomassie Brilliant Blue staining.

### Glucose depletion and the induction of autophagy. Autophagy induced by prolonged starvation precedes the development of programmed cell death

To demonstrate that metabolic stress (through glucose depletion) induces autophagy that correlated with protein hydrolysis, monodansylcadaverine (MDC), a common autophagosome marker, was used to study the development of autophagy in BY-2 cells (Biederbick A, Kern HF, 1995). At the morphological level, autophagic processes were characterised by formation of multiple vacuolar structures localised diffusely in the cytoplasm or formation of clusters in perinuclear space (Fig. 2A). Microscopic and colorimetric studies revealed a well-defined time synchronisation of glucose depletion in culture medium, lower concentrations of cellular proteins, and an increase in the percentage of cells with morphological features of autophagy. Thus, the percentage of autophagic cells did not exceed 3% with the presence of sugars in the medium in initial stages of cultivation. Sharp lowering of glucose concentrations from day 7 on led to an increase of the proportion of MDC-positive cells of up to 7%. After further development of this process, the quantities of cells containing autophagosomes amounted to 15%, 19%, and 31% of the total number of cells on days 8, 9, and 10 of cultivation, respectively. This measure attained its maximum level of 42% on day 11 (Fig. 2B).

**Fig. 2.**
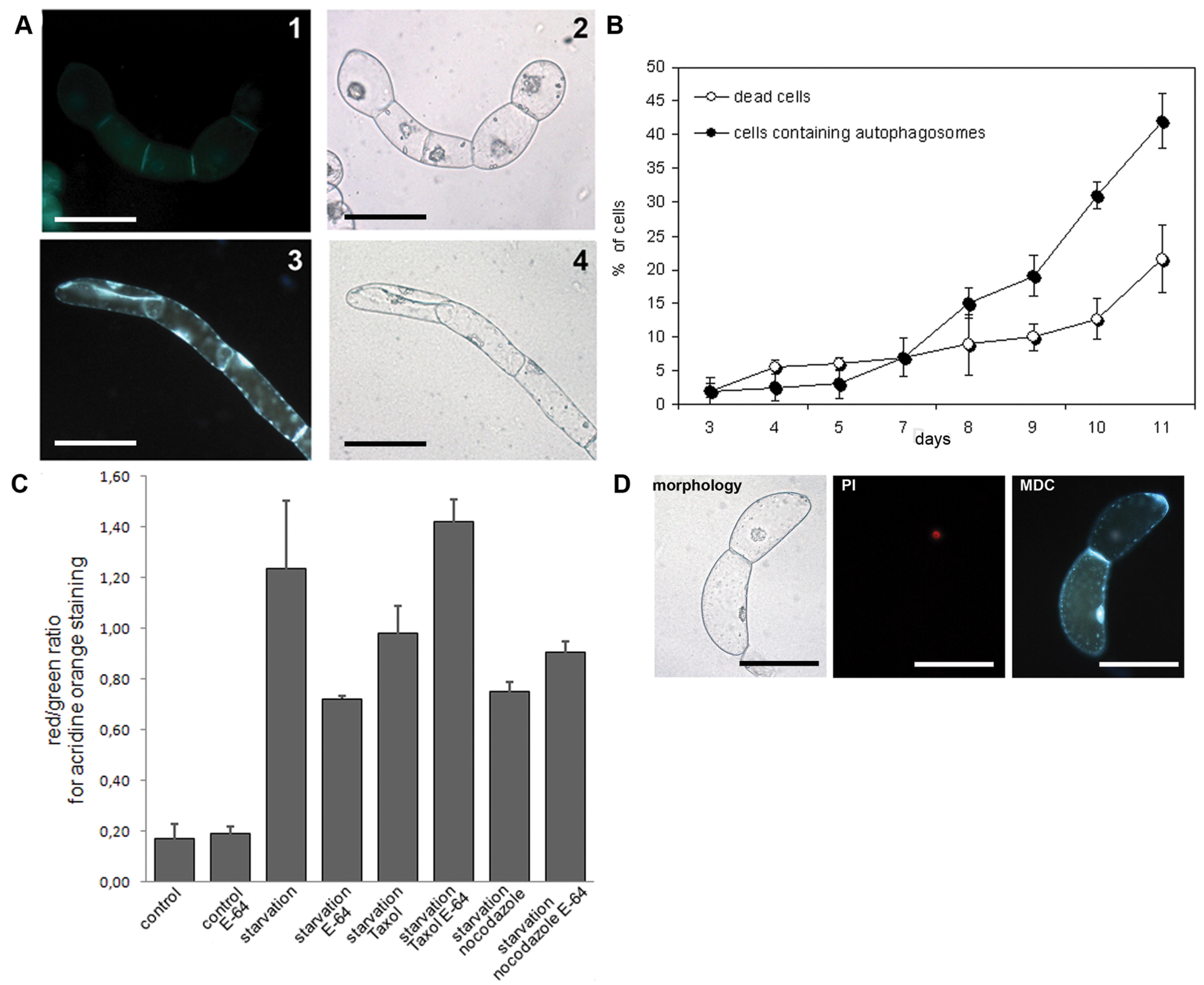
Autophagy development in BY-2 cells under cultivation scheme 1. (A) Morphologic features of autophagy; 1,2 – cultivation day 4, characterised by lack of autophagy. 3,4 – cultivation day 9, characterised by cells with multiple autophagosomes. 1,3 – MDC staining. 2,4 – general morphology. (B) Dynamics of cell survival rate and autophagy development. (C) Confocal measurements of cytoplasm acidification levels in proliferating and starving cells in combination with E-64, taxol and nocodazole treatment; the presented values are mean red/green AO fluorescence ratio (100 cells of tree independent experiments were measured). (D) Cell death can be a consequence of autophagy (upper cell is dead but contains multiple autophagosomes) (PI/MDC staining, day 9). Bar 50 μm.

The above evidence suggests that development of autophagy is an adaptive trophic mechanism under metabolic stress, one aspect of which is the energy source depletion in culture medium. It should be noted that BY-2 cells had a relatively stable survival rate (determined using propidium iodide staining) under long-term cultivation. The standard cultivation period was characterised by low lethality that did not exceed 7%, whereas the cell death rate was 21.5% on day 11 under prolonged cultivation. A high cell survival rate indicates the reliability of the obtained results and adequacy of the used starvation model to reproduce the autophagic conditions in the BY-2 cell line. Moreover, this tendency persisted when cells were cultivated for 15 days in cultivation scheme 2. A rapid increase of cells containing autophagosomes occurred from day 8 to day 11 (24 to 27% of cells, respectively) that corresponded with sucrose exhaustion and the start of protein hydrolysis. Further development of cellular events on subsequent days, however, was more significant. On day 14, the number of cells containing autophagosomes decreased rapidly (14%); on day 15, such cells were not present at all. However, in this time range we detected a clear inverse correlation between the autophagy progress and programmed cell death development. Thus, the amount of cells with nucleosomal DNA fragmentation was at a constant level between 2.5 and 3.1% against a background of culture growth and autophagy development throughout the whole cultivation period from days 1 to 14. However, on days 14 and 15 the quantity of terminal dUTP nick end labeling (TUNEL)-positive cells increased rapidly, reaching 4.8% and 27%, respectively (Fig. 4). Additionally, inhibition of autolysosome formation by cystein inhibitor E-64 was used to prove reliability of autophagic flow. E-64 was added in the culture medium aseptically at the termination of exponential growth stage (5 or 7 day dependent of experimental scheme). In both cases dramatic development of cell death was detected by PI staining (not shown). Autophagy inhibition under metabolic stress was resulted in near total mortality of the culture till 9 and 11 days, correspondingly. Under cultivation scheme 1 this effect was much less pronounced, probably because of the faster development of cell depletion (not shown).

**Fig. 3.**
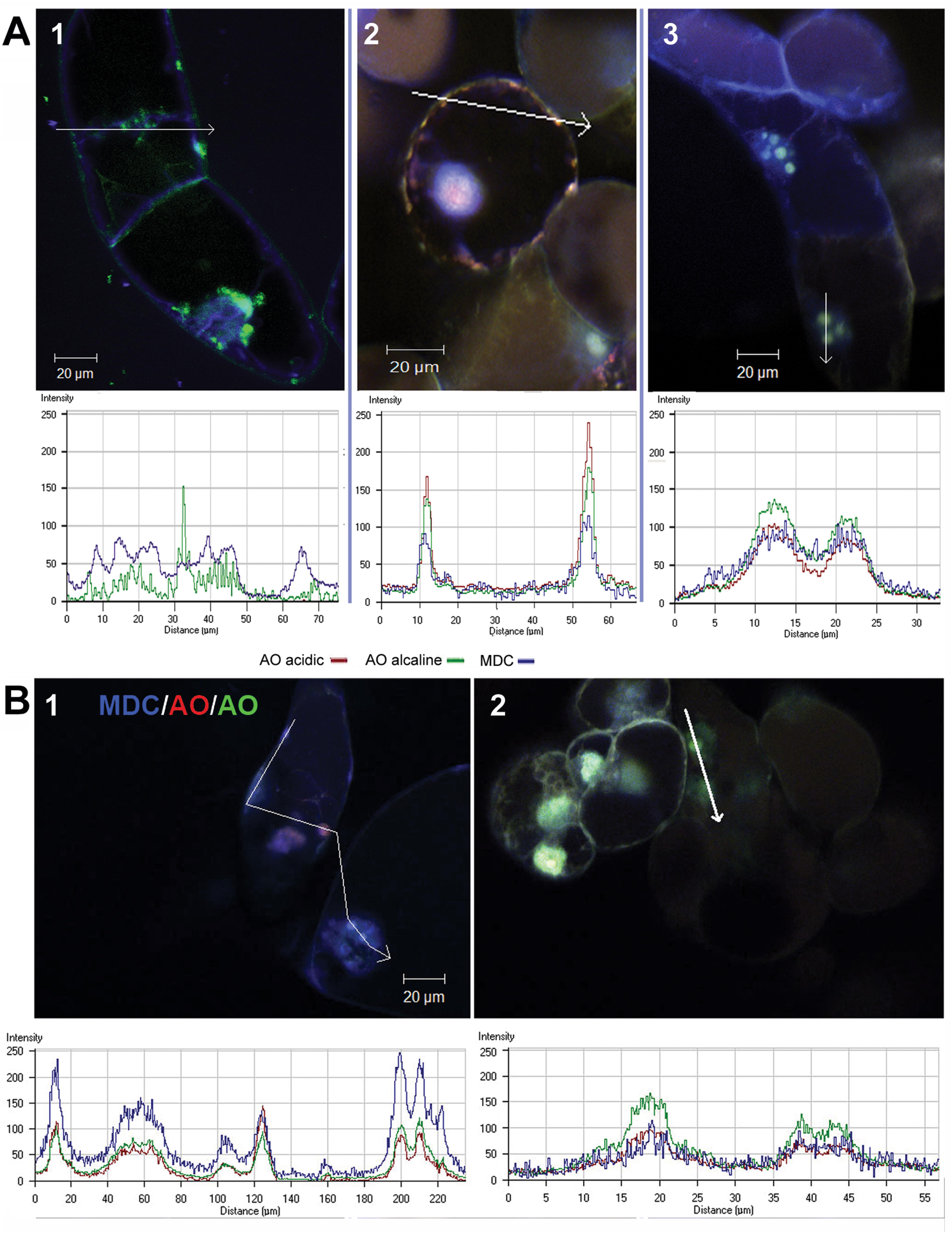
CLSM investigation of cellular autophagosomes distribution and cytoplasm acidification under autophagy inhibition (A) and taxol/nocodazole treatment (B). A1 – control cells in exponential growth stage, A2 – starved cells in autophagy development stage, A3 – starved cells pretreated with E-64 in autophagy development stage, B1 and B2 – tarved cells in autophagy development stage pretreated with taxol and nocodazole, correspondingly. AO/MDC staining.

**Fig. 4.**
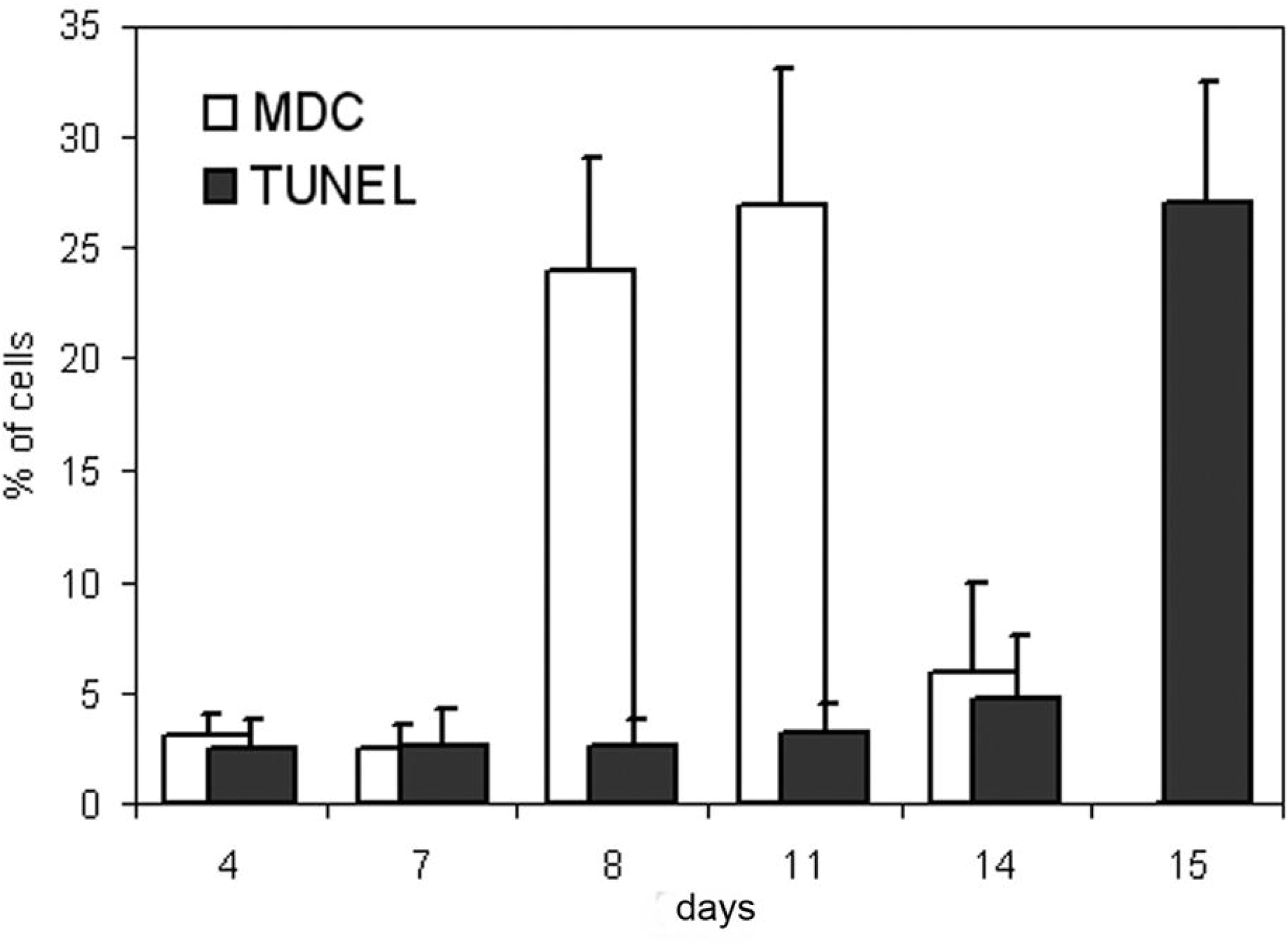
Development of autophagy and nucleosomal DNA fragmentation in BY-2 cells after 15 days of cultivation. MDC – cells containing autophagosomes stained by monodansylcadaverine. TUNEL – TUNEL positive cells.

### Synergistic impact of starvation, inhibition of autophagy and microtubular drags on the cytoplasm acidification and intracellular autophagosome flow

Additionally, it should be noted that cell acidification is among the morphological hallmarks that relate to programmed cell death and would be specific to both cultivation schemes. This characteristic (determined by acridine orange staining (Zelenin, 1966)) was observed as a relatively massive phenomenon in the cells at the stage of transition to autophagy and the appearance of protein hydrolysis. It is important that acidification was observed mainly in the cytoplasm and vacuoles, but not in the nuclei where it was a rare event. En masse, acidification level in starving cells (exponential growth stage) was in near seven fold higher than in control cells (autophagy development stage; additionally to that inhibition of autophagy and treatment with microtubular drugs decreased acidification level under starvation conditions (Fig. 2C). On the other hand, we could observe cell death as a co-occurring result of autophagy in both experimental schemes (Fig. 2D). This observation supports two arguments; namely, that autophagy and apoptosis are two discrete adaptive processes (under similar stress conditions) and autophagy should be regarded as one of the stages of programmed cell death. Both cultivation schemes showed quite similar characteristics of the development of studied processes (coincidence of glucose depletion, protein hydrolysis and autophagy induction). However, scheme 2 was mainly used for biochemical studies of the functional state of the cytoskeleton due to less dynamic development of cell death in the final cultivation stages of the experiments.

Inhibition of autophagy during metabolic stress leads to interruption of intracellular autophagosomes traffic and their localization in the perinuclear space. Notably, this effect was also observed when the action of taxol and nocodazole in starving cells. Both observations indicate microtubules involvment in intracellular transport of autophagosomes and mediation of autophagy (Fig. 3).

### Development of autophagy and *α*-tubulin acetylation and detyrosination

The role of tubulin acetylation on Lys40 as a regulatory modification is critical both in the development of starvation-induced autophagy (Xie et al., 2010; Geeraert et al., 2010) as well as in the neurodegenerative processes mediated by autophagy (Perez et al., 2009). The dynamics of tubulin acetylation and detyrosination was investigated in the development of stress-induced autophagy in plant cells by considering the role of acetylation in the attracting of motor proteins to microtubules (Reed et al., 2006) and the potential impact of detyrosination in these processes (Konishi and Setou, 2009; Hammond et al., 2010; Walter et al., 2012).

Generally, a clear correlation was present between the dynamics of glucose depletion and tubulin acetylation. Samples obtained by cultivation scheme 2 were used for biochemical investigations of the MTs functional state. First, increases in the degree of this modification were detected in the time interval that defines the end-stage of the stationary growth phase and is the starting point of culture exhaustion (day 7). Furthermore, the acetylation level becomes maximal at the time interval that corresponds to dramatic changes in glucose concentration and to the highest values of autophagy. This process is reversible and reduction of tubulin acetylation synchronously precedes the attenuation of autophagy in BY-2 cells: the maximum level of modified tubulin on days 8 and 9 declines on day 10 and corresponds to the maximum level of autophagy on days 8 through 10 (24-27% of MDC positive cells) (Fig. 5A,B). In turn, basal autophagic levels were accompanied by lack of tubulin acetylation at the terminal experimental points. At the same time, the dynamics of gains in tubulin detyrosination was slightly different. Although the first signs of this modification were observed on day 8, the highest values for tubulin detyrosination were reached from day 9 to day 11. This post-translational modification of tubulin does not match in time with acetylation completely under the described experimental conditions. Based on the behaviour of the experimental dynamics, it can be argued that tubulin detyrosination is related to stress-induced autophagy as well as to growth phase stages in cells BY-2 and is not correlated (or is weakly correlated) with changes in glucose concentration. In general, similar dynamic behaviour for both modifications was observed for the 11-day cultivation scheme 1 (not shown). An additional important fact of note is the decrease of tubulin acetylation and detyrosination levels in the development of nucleosomal fragmentation in the 15-day cultivation scheme.

**Fig. 5.**
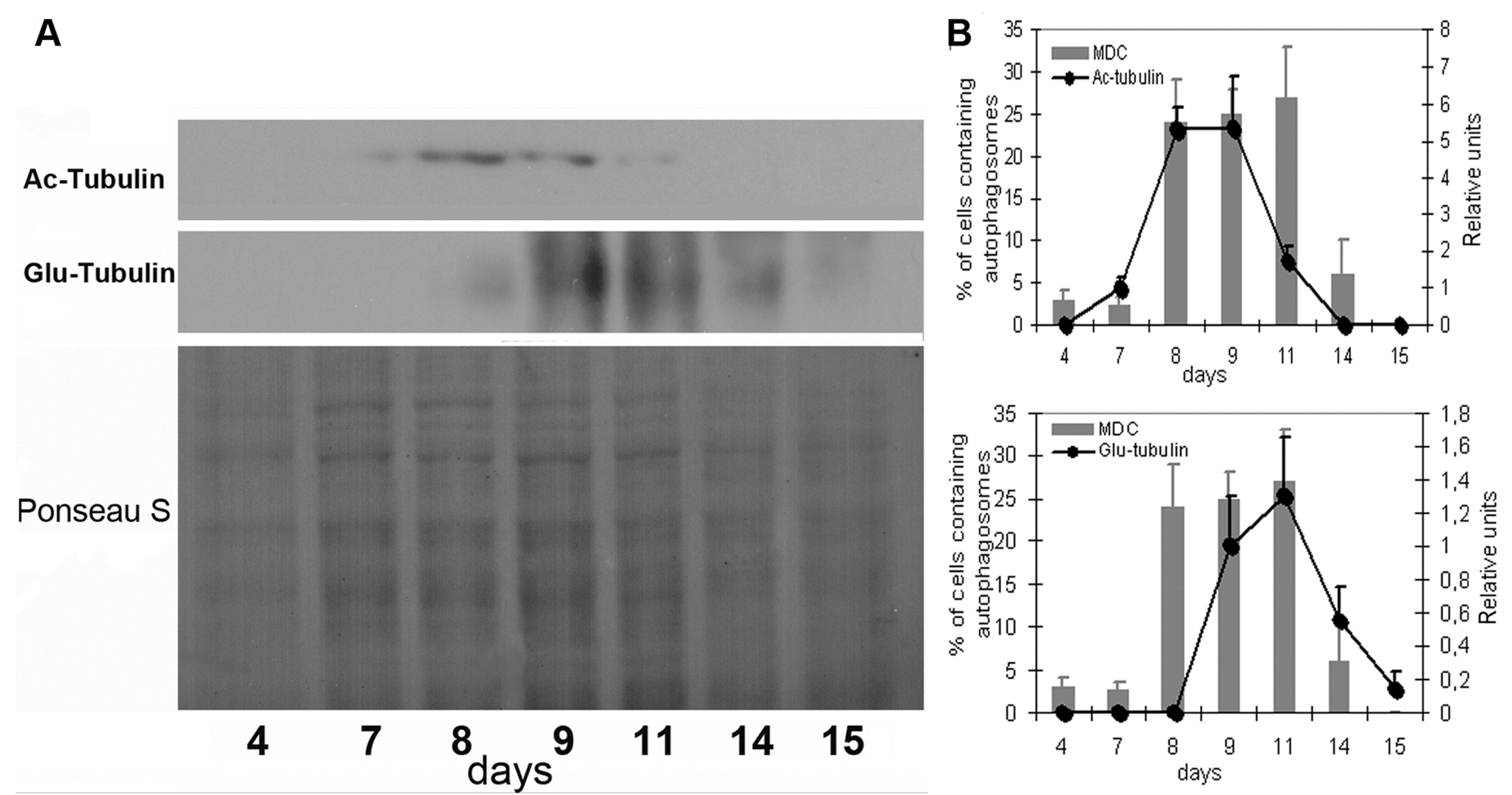
Acetylation and detyrosination of *α*-tubulin under autophagy development in BY-2 cells (starvation scheme 2). (A) Western blot analysis of tubulin modifications. (B) Averaged ratio of the amount of cells with autophagy and tubulin modification levels.

### Starvation-induced post-translational modifications of *α*-tubulin and their molecular microenvironment changes

We performed Co-IP with subsequent analysis of the protein complexes using the Experion system (Bio-Rad Laboratories; Hercules, CA, United States) to evaluate the affinity of modified tubulins to cytosolic proteins and to compare protein pools related to microtubules under conditions of autophagy.

Three reference time points of cultivation scheme 2 were chose to reflect different stages of culture growth (day 4 – exponential growth stage; day 11 – autophagy development; day 14 – transition from autophagy to apoptosis). As a result, signs were found indicating the crucial influence of post-translational modifications of tubulin (such as acetylation and detyrosination) on the microenvironment of MTs. Moreover, the samples of varied time points precipitated by the same antibodies were significantly different by composition of Co-IP complexes. It should be noted that content of tubulin-like fractions in the comparison groups (samples from days 4, 11 and 14) were virtually unchanged. The sets of precipitated proteins that have molecular weights in the range of 45-58 kDa were accepted as α-tubulin, despite the need to use this criterion with some caution.

In the Co-IP procedure, BY-2 cell lysates were incubated with 10 μg of antibodies immobilised on coupling resin; a significant excess of lysate in reaction mixture was used to ensure relative uniformity of antigen binding (tubulin and its modified forms). This uniformity was generally achieved at all experimental time points. Mean concentration levels for tubulin-like proteins in analysed samples were finally quantified via Experion system analysis within a range of 13-29 ng μl^−1^. Extremely low concentrations of the bait protein were an exception, only occurring in one case (in the precipitate of antibodies against acetylated tubulin from the sample of 14 2.61 ng μl^−1^). This value is located on the lower border of validity of the method and shows the disappearance of this modification with the fading of autophagic signs. The differences among the precipitated proteins of 45-58 kDa range were generally minimal within the same precipitating antibody. Shifts in tubulin molecular weights and certain individual differences within experimental groups can be explained by the presence of other post-translational modifications that could alter electrophoretic mobility (Fig. 6A,B; Table 1).

**Fig. 6.**
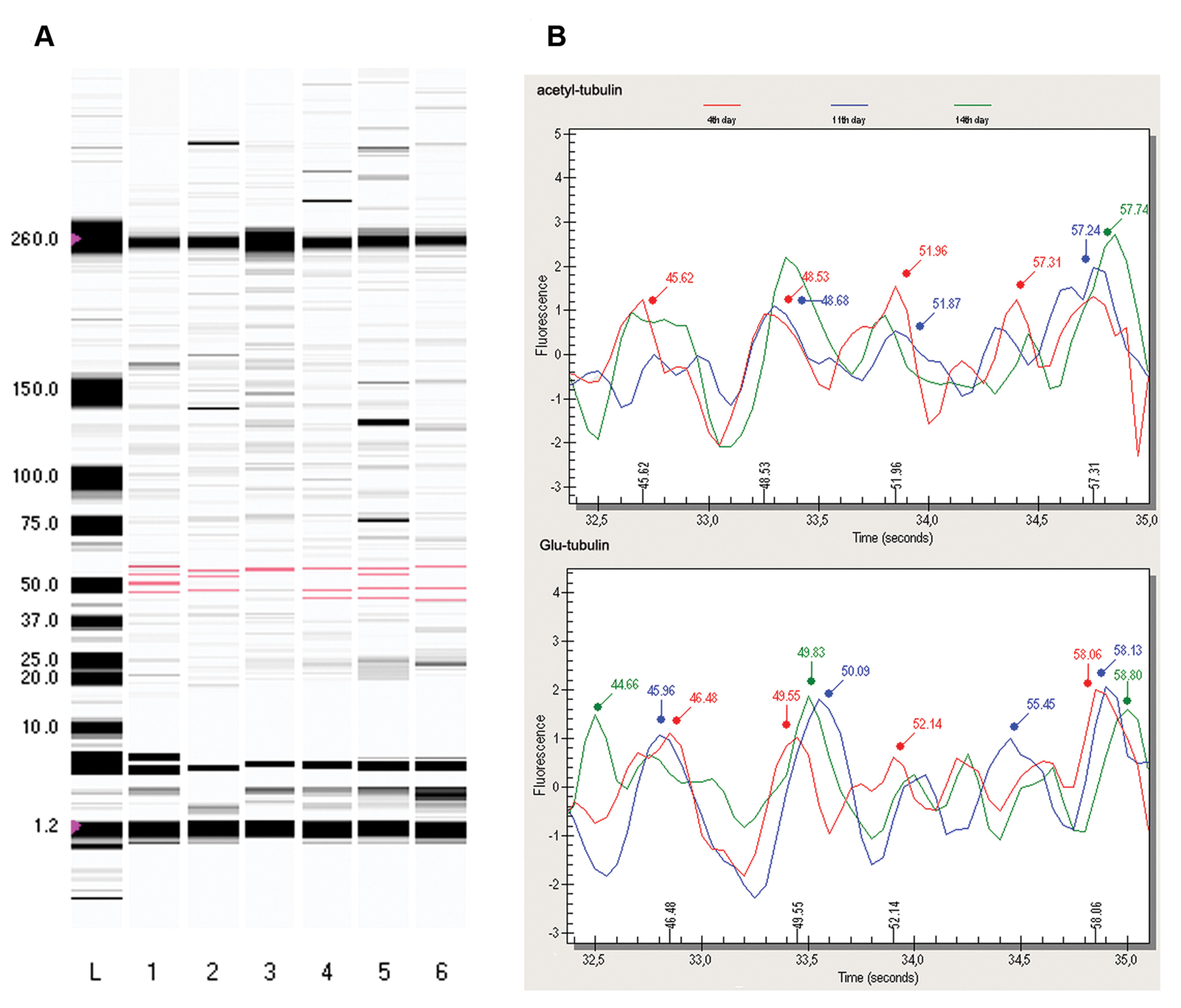
Modified tubulins from Co-IP samples determined by Experion analysis. (A) Digital gel, lanes from 1 to 6: 1-3 – precipitation by anti-acetylated tubulin antibodies; 4-5 – precipitation by anti-Glu-tubulin antibodies; 1 and 4 – day 4 samples, 2 and 5 – day 11 samples, 3 and 6 – day 14 samples (bands corresponding to tubulins; molecular weights colored in red). (B) Densitometric graph of precipitated tubulin-like proteins.

**Table 1.**
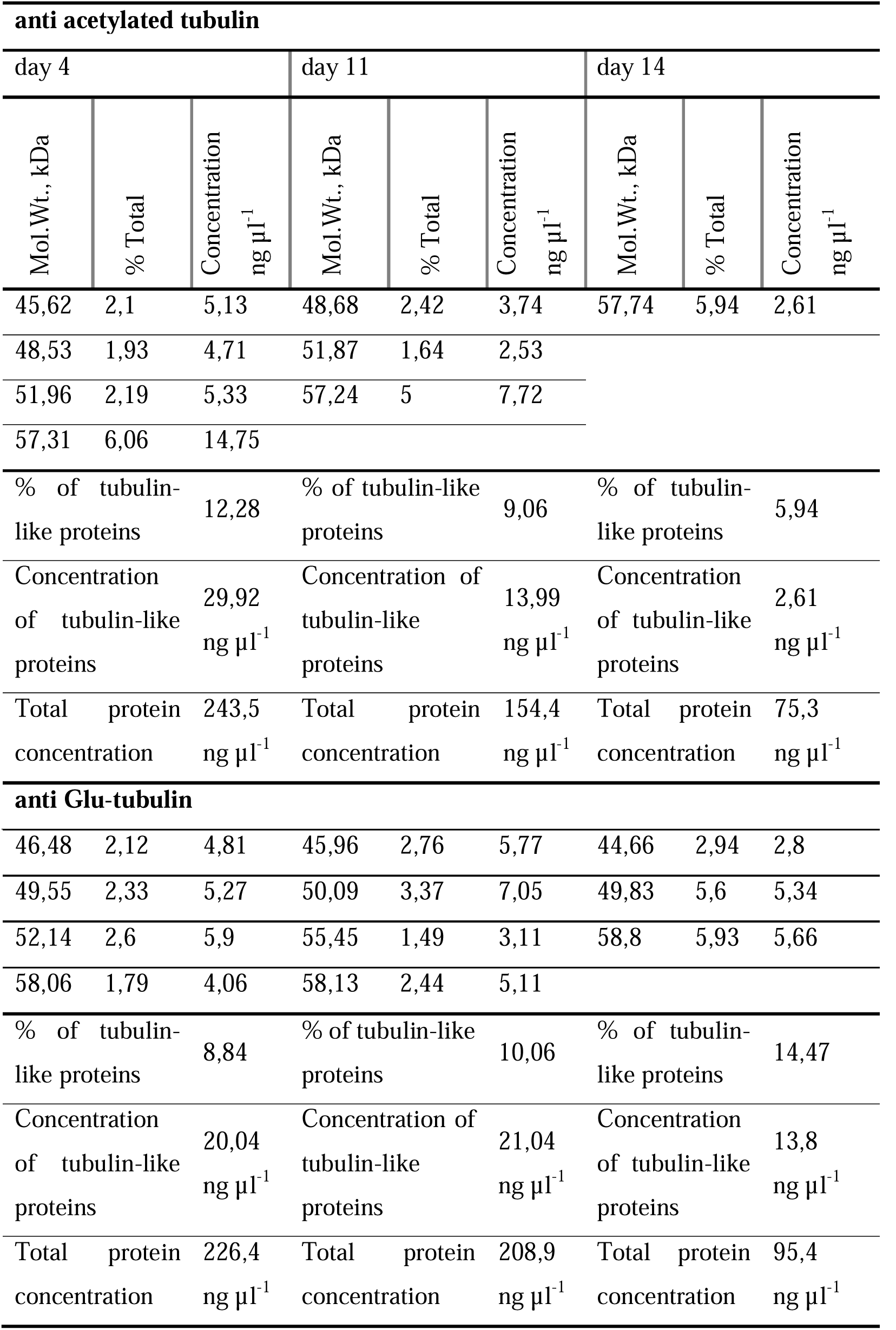
Quantitative data of precipitated tubulin-like fractions.

By co-precipitate analysis, we considered prey-proteins as valid when their values were not less than 4.5 % of the total protein amount. Comparison of bait:prey complexes revealed significant differences in the microenvironments of acetylated and detyrosinated tubulin as well as among varied time points within the same precipitating antibody. Certain co-precipitated proteins could be associated with the exponential growth phase and stage of autophagy (4 and 11 days), the other proteins for the stage of autophagy and time point that was characterised by signs of programmed cell death (14 and 15 days). In particular, this tendency was observed in the complexes of acetylated tubulin; the first (day 4) and second (day 11) experimental points revealed coincidences in the spectrum of small and medium proteins, while the second and third experimental points (day 11 and day 14) possessed some similarity to high-molecular weight proteins. The detyrosinated tubulin microenvironment showed dramatic differences in protein composition compared to the acetylated tubulin complexes. On the one hand, the microenvironment was characterised by proteins that persisted during all experimental stages; on the other hand, some proteins were detected occasionally in the initial or the final stage of the experiment (Table 2).

**Table 2.**
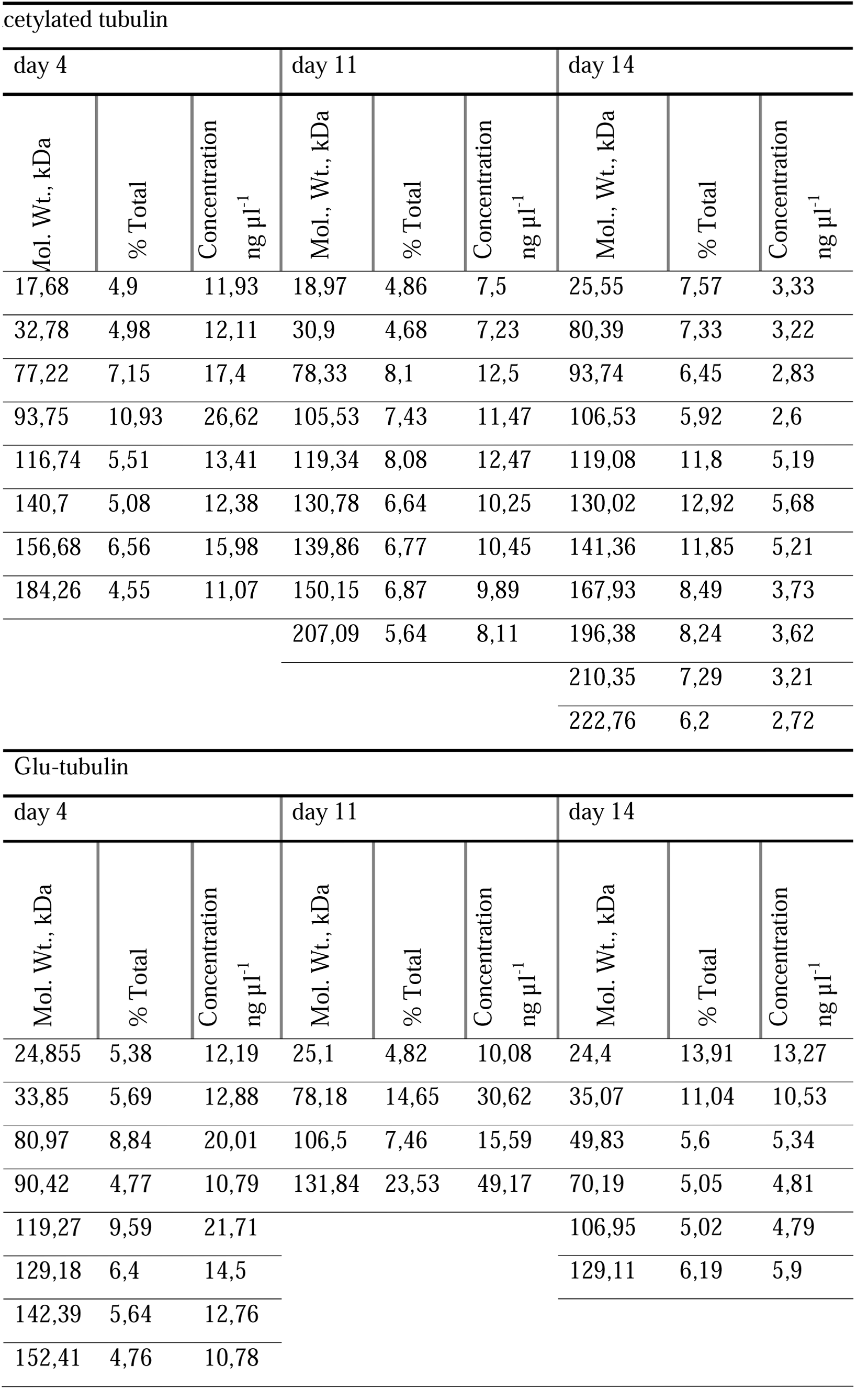
Protein spectrum that forms a complex with modified α-tubulin from the samples that correspond to different growth stages (cultivation days 4, 11 and 14).

Significant differences of prey-proteins that form complexes with modified tubulin indicate the regulatory role of these modifications when supporting the functional state of the cytoskeleton under trophic stress conditions. Thus, it should be selected following prey-proteins that can potentially be involved in the mediation of autophagy by cytoskeleton (detected in BY-2 cell samples on day 11): 78.3; 105.5; 119.3; 130.8; 139.9; 150.2 kDa co-precipitated by antibodies against acetylated tubulin and 25.1; 78.2; 106.5; 131.8 kDa co-precipitated by antibodies against detyrosinated tubulin (Table 2). Unexpectedly, we did not identify proteins corresponding to tobacco homologues of Atg8 with predicted molecular weight in the range of 14 kDa in any of the samples. This fact may indicate an indirect interaction of tubulin and Atg8 and implementation of other MAP proteins in the formation of this complex (Mann et al., 1994; Xie et al., 2011).

## DISCUSSION

There is no doubt that certain nutrients are essential for development of degradation processes in plants under different physiologic and stress conditions. For the concept, carbohydrates hold a special influence in that they act not only as a source of carbon but also perform regulatory functions in cell signalling. Thus, several basic mechanisms of glucose reception and signalling that affect the regulation of gene expression as well as a significant number of cellular processes were discovered in plant cells: namely, hexokinase (HXK)- dependent, HXK-independent and glycolysis-related pathways (Xiao et al., 2000; Rolland et al., 2006). Autophagy as a consequence of sucrose depletion was widely shown in plants previously (Chen et al., 1994; Contento et al., 2004; Xiong et al., 2005; Bassham et al., 2006; Rose et al., 2006; Wingler et al., 2009; Honig et al., 2012). A very promising potential link between glucose signalling and autophagy induction is SNF1-related serine/threonine-protein kinase SnRK (sucrose non-fermenting-1-related protein kinase-1) that regulates a signal transduction cascade of carbohydrate metabolism in higher plants (Halford et al., 2003). Notably, it was clearly shown that overexpression of its catalytic subunit (KIN10) in Arabidopsis seedlings (induced by sucrose starvation) led to elevated expression of ATG genes (Baena-González et al., 2007). On the other hand, KIN10 is a plant orthologue of the catalytic subunit of mammalian AMP-activated protein kinase (AMPK) that negatively regulates activity of the mTOR signaling pathway (Kim et al., 2011). So, whereas mTOR kinase signalling regulates plant autophagy development (Liu and Bassham, 2010; Xiong and Sheen, 2014) and regulation of TOR activity is glucose-dependent (Xiong and Sheen, 2012) we presumed a significant regulatory impact of dynamic sugar depletion on the induction of autophagy in our experiments. For these reasons, a long-term cultivation scheme was chosen as a good way to track the dynamic flow of autophagy (basal and starvation-induced) as well as programmed cell death under a background of metabolic stress. The data obtained by 11- and 15-day cultivation of BY-2 cells indicate that a crucial glucose level in the culture medium is an initial trigger that starts the process of autophagy. For both experimental schemes, we observed a clear time synchronisation of three events: glucose depletion, protein hydrolysis and an increase in the number of cells with features of autophagy. It should be noted that there was some basal level of autophagy (less than 3% of cells) that was characteristic of the early cultivation stages in the absence of nutritional deficiency. This in turn allowed us to predict the presence of a discrete regulatory mechanism of basal (physiological) autophagy in plant cells that is independent of glucose signalling. We can consider this cellular event as a housekeeping process for molecular/organelle turnover or vacuole formation (Inoue et al., 2006; Yano et al., 2007). But since the level of glucose is declining (day 7 and day 9 of 11- and 15-day cultivation, respectively), it is clearly seen that protein degradation and autophagy develop hand in hand (Fig. 1-2).

An entire picture of cell death processes developing under prolonged depletion of nutrients is rather complicated. On one hand, only under slow starvation conditions (15 days) did a reliable apoptosis-like cell death develop, the levels of which rapidly increased during the terminal stage (day 15) and which was characterised by nucleosomal DNA fragmentation (Fig. 4). On the other hand, we have seen some small percentage of TUNEL-positive cells at all stages of culture growth, indicating the presence of an apoptosis-like process that is a largely physiologic and stress-independent phenomenon. Moreover, under 11-day cultivation conditions we did not detect the same type of programmed cell death as a massive final event in spite of the presence of the aforementioned cells with nucleosomal DNA fragmentation on the basal level (not shown). In both cases we observed a similar number of dead cells at the end of experiments (21.5% and 27%, respectively, at 11 and 15 days); however, these cells died in different ways. Indeed, beginning from the stage of exponential development of autophagy, the majority of experimental points for both experimental schemes were characterised by the presence of cells that were dead but contained autophagosomes (Fig. 2D). Thus, it is possible to predict cell death as one of the possible scenarios of autophagy development. Autophagy may play a role not only as a survival mechanism, but also as a ‘pro-death’ process in animal and plant cells under certain conditions (Tsujimoto and Shimizu, 2005; Patel et al., 2006; Cacas and Diamond, 2009). Autophagy-like and apoptosis-like processes can be involved both during implementation of plant development (formation of root and vascular tissues) as well as in response to external stimuli. Although autophagy-like cellular events are mostly associated with the aging process, response to abiotic factors and involvement in innate immunity (Liu et al., 2005; Love et al., 2008), it is impossible to track the synchronicity between autophagy and nucleosomal DNA fragmentation in individual cells under our experimental conditions. On the other hand, despite the generally accepted understanding of the regulation of autophagy and apoptosis in this case we can’t assert that DNA fragmentation is not a consequence of autophagy, as it was observed in death of developmental nurse cells in the *Drosophila melanogaster* ovary, in which autophagy acts as a trigger of cell death (Bass et al., 2010; Nezis et al., 2010). The results obtained during this study provide us the reasons to anticipate a simultaneous flow of programmed cell death and autophagy, both of which are probably regulated by the same set of external stimuli and compete with each other. It was suggested that autophagy-like cell death acts antagonistically to apoptosis-like programmed cell death and furthermore that signalling pathways of jasmonic acid, salicylic acid, and reactive oxygen species (ROS) as well as ethylene are general for regulation of both cell death scenarios. However, triggering and overlapping between death signalling pathways are in large measure of a temporal nature (rapid apoptotic cell death, slow autophagic cell death) (Love et al., 2008) that conforms with our results.

Acidification of the cytoplasm was another unifying feature that was observed under different levels of DNA fragmentation (Fig. 2C). This phenomenon had clear temporal relationship between arising of cytoplasm acidification, medium depletion, protein hydrolysis and development of autophagy. Given that cytoplasm acidification (the outcome of malfunction of intracellular proton export systems) is one of the hallmarks of apoptotic cell death (Li and Eastman, 1995; Gottlieb et al., 1995; Boyle et al., 1997), it is possible to predict the development of such cellular scenarios in BY-2 cells under metabolic stress conditions.

So, further research is required into how switching between pro-survival function of autophagy and the ‘pro-death’ function occurs. In other words, given a background of metabolic stress, in which cellular events does apoptosis start to prevail over autophagy? We can speculate on the induction of apoptosis by ROS accumulation in the final stage of medium exhaustion that is able to regulate the development of apoptotic death. The mechanisms that synchronise apoptotic and autophagic signalling cascades in time are unclear as yet. We can also provide a regulatory role of some secondary metabolites that are accumulated in the medium in the later cultivation stages, certain concentrations of which can be the impetus for PCD development.

The cellular events described in this study took place against the backdrop of significant changes in the functional state of the cytoskeleton. Namely, these changes included changes in the acetylation and detyrosination level of tubulin and the associated variability of the cytoskeletal proteins microenvironment. The most important pattern was temporal relatedness of these modifications with the development of autophagy and apoptosis-like events. A certain low basal level of tubulin acetylation can be observed in cells in the exponential growth phase, indicating that the role of this modification in plants is broader than simply being a response to pathological conditions. This modification has been shown for numerous plant species, including *Nicotiana tabacum* (Smertenko et al., 1997), *Arabidopsis thaliana* (Tran et al., 2012), *Zea mays*, *Triticum turgidum*, *Vigna sinensis* (Giannoutsou et al., 2012) as well as for other monocots and dicots (Nakagawa et al., 2013*b*). It was determined that tubulin acetylation is dynamically regulated during development of plant organs and this modification is tissue-specific (Wang et al., 2004; Nakagawa et al., 2013a,b). In plants, tubulin acetylation has been shown to be an inherent modification for all types of microtubular arrays, including interphase cortical MTs, preprophase band, mitotic spindle and phragmoplast (Smertenko et al., 1997). This modification was demonstrated to be associated with cortical endoplasmatic reticulum by preprophase band formation in dividing cells of maize, wheat and cowpea (Giannoutsou et al., 2012) and reflects the stable state of MTs (Huang and Lloyd, 1999; Giannoutsou et al., 2012).

Detyrosination is a conserved post-translational modification that occurs by tyrosine removal on the C-terminus of polymerised α-tubulin catalysed by unidentified carboxypeptidase (Verhey and Gaertig, 2007). It is known that detyrosinated and tyrosinated MTs are able to interact differentially with two types of microtubule-binding proteins. Molecular motors, namely kinesin-1, participate in cellular cargo movement along detyrosinated MT, but tyrosinated one recruits plus-end tracking proteins (+TIPs) containing a CAP-Gly domain (Hammond et al., 2009). On the other hand, tubulin detyrosination can also reflect the MT stable state (Khawaja and Gregg, 1988) as well as mediate processes of cellular differentiation, growth and motility (Gundersen and Bulinski, 1988; Rodionov et al., 1994). The functions of this modification require further investigation in plants.

According to our results, the appearance and disappearance of tubulin acetylation are completely synchronised in time with the development and termination of autophagy in BY-2 cells during metabolic stress. These data are in good agreement with possible functions of this modification in the development of autophagy in mammals. Thus, starvation-induced tubulin hyperacetylation in HeLa cells leads to recruitment of kinesin-1 with subsequent Jun amino-terminal kinases (JNK) activation that results in induction of MT-associated autophagosome formation (Geeraert et al., 2010). Control of autophagosome formation and movement depends on compartmentalization between dynamic and stable MTs and is regulated by molecular motor activities. But processes of autophagosome formation are probably more related to dynamic acetylated MTs, whereas autophagosome traffic concerns stable ones (Mackeh et al., 2013). It is interesting that from another point of view the JNK signalling pathway might be linked with autophagic death regulation under Bax/Bak-deficient background (proapoptotic members of the Bcl-2 family) (Tsujimoto and Shimizu, 2005; Shimizu et al., 2010). But autophagy that promotes cell survival does not depend on JNK (Tsujimoto and Shimizu, 2005). Some shorter accounts of these processes complicate the argument concerning the role of detyrosination in autophagy. But the phenomenon of the co-existence of both modifications during the aforementioned processes provides a reason to consider detyrosination as an autophagy-related modification. It may reflects the similar processes as selective kinesin-1 traffic in detyrosinated MTs (Dunn et al., 2008; Konishi and Setou, 2009) or be bolstered with the evidence that both tubulin acetylation and detyrosination are involved in supporting the stable state of MT (Cambray-Deakin and Burgoyne, 1987). Most likely, a relationship exists between glucose signalling and regulation of tubulin acetylation, because medium depletion clearly coincides in time with increasing modification levels. Increases of tubulin detyrosination are slightly behind the development of autophagy, which could indicate an indirect involvement of this modification in implementation of autophagy but also the development of processes similar to aging mechanisms.

In addition to the above we can confidently assert that apoptosis-like processes (nucleosomal DNA fragmentation) and acetylation/detyrosination of tubulin are mutually opposite cellular events (Fig. 4,5). On the other hand, at the time when the apoptotic scenario becomes apparent, reduction of the detyrosination level is likely to indirectly indicate the increase of tubulin nitrotyrosination, which, in turn, may denote a disruption of MTs (Yemets et al., 2011). The data also strongly suggest that tubulin acetylation and detyrosination may have a reversible nature (particularly in the processes associated with programmed cell death) and are dependent on the functional state of plant cells under metabolic stress conditions (Fig. 5). Taking into account the data that tubulin tyrosine ligase sequences exist only in mammalian and trypanosomes genomes (Janke et al., 2005) it is possible to predict the existence of plants’ TTL-like enzymes to provide tyrosination/detyrosination cycling of plant tubulin. At the same time, α-tubulin deacetylation is probably catalysed by histone deacetylase 14 (HDA14), as shown in *Arabidopsis thaliana* (Tran et al., 2012). Interestingly, histone deacetylase 6 (HDAC6) (that is able to deacetylate human α-tubulin (Hammond et al., 2009)) appears as a master regulator of the cell protective response to cytotoxic protein aggregate formation in mice cells (Boyault et al., 2007). In addition, we had shown some basal levels of both modifications during the initial stages of the experiments, indicating the physiological function of these modifications under normal conditions.

The comparative picture of the microenvironment of acetylated and detyrosinated tubulin is quite dissimilar and coincides only partially in most cases (Fig. 6A, Table 2). These findings allow us to assume the possibility of simultaneous existence of both modifications on the same molecule as well as on individual tubulin molecules. The presence of high molecular weight proteins (more than 105 kDa) in precipitated samples gives us some reason to expect tobacco kinesin-1 homologues in these complexes and is consistent with the above mentioned data related to the role of these modifications in conjunction of kinesin-1 and MT. There is some evidence of kinesin-1-like proteins in tobacco cells despite the significant differences of kinesin superfamily in plants compared to other eukaryotes(Reddy and Day, 2001). Thus for example, kinesin-like proteins NACK1 (108,6 kDa, UniProt number – Q8S950) and NACK2 (107,2 kDa, UniProt number – Q8S950) regulate activity and phragmoplast localisation of mitogen-activated protein kinase NPK1 in BY-2 cells (Nishihama *et al.*, 2002). Other TKRP125 (125 kDa kinesin-related protein, 113,7 kDa, UniProt number – O23826) exhibits its involvement in the cell cycle-dependent changes in MT arrays, including the organization of the phragmoplast, and in the movement of chromosomes in anaphase cells (Asada and Shibaoka, 1994; Nishihama et al., 2002). But statements about involvement of kinesin-like as well as other proteins to MT-mediated process of autophagy requires further investigation in plant cells.

## MATERIAL AND METHODS

### Cell culture and starvation conditions

BY-2 tobacco cells were cultivated under standard conditions (Nagata et al., 1992) in 100 ml of liquid MS medium in 250 mL Erlenmeyer flask at 110 rpm using a Heidolph Unimax 1010 rotary shaker equipped with Heidolph Incubator 1000 at 26°C in the dark (Heidolph North America; Elk Grove, IL, United States). Sub-cultivations were performed by passaging of 2 ml of cell suspension that corresponded to the near-overgrown stage every 7 days. In starvation experiments, the duration of cultivation time was extended up to 11 and 15 days to induce autophagy (called starvation scheme 1 and 2, respectively). Maximal cell survival in the end-stage of the experiment was the main criterion for determining the duration of cultivation time (no more than 30% of dead cells). The cell suspension volume was similar with sub-cultivation volume for passaging by 11 days cultivation (scheme 1), but for the 15-day cultivation scheme (scheme 2) we decreased the inoculation volume up to 0.5 ml.

One ml of cell suspension was collected aseptically to measure cellular proteins and glucose levels as well as to perform microscopic assays on days 3 through 5, 7, 8, 10, 11, 14 and 15. Samples for the TUNEL assay were collected on days 4, 7, 8, 11, 14 and 15. Finally, cell suspensions of 1 ml were sedimented at 500 g for 2 min and washed once with phosphate buffered saline (PBS) on days 4, 7, 9, 11, 14, and 15. Collected cells were stored in liquid nitrogen for further biochemical analysis.

### Autophagy inhibition, application of MT targeting drugs

To keep the culture from autophagy development specific permanent inhibitor of lysosomal serine protease E-64 was used (78434, Thermo Fisher Scientific Inc., Waltham, MA, United States). Sterile inhibitor stock solution in ethanol was added to the culture to final 100 μM concentration on 5 and 7 days for experimental scheme 1 and 2, respectively.

To investigate interrelation between autophagy development and microtubules simultaneous cell treatments with E-64, taxol (microtubule stabilizing agent) and nocodazole (microtubule depolymerized agent) were used. Cells were pretreated (or not) with E-64 within 24 h and, then, subjected to 10 μM taxol or 1 μM nocodazole treatment within 3 h. After that cells were analyzed immediately by CLSM to determine autophagosome formation and cellular distribution as well as acidification level of cytoplasm.

### Dye loading and TUNEL assay

For microscopic estimation of cell viability, morphology and autophagy development the following dyes and dilutions were used: propidium iodide (PI) – 1.5 μg ml^-1^ (81845, Fluka); 4,6- diamidino-2-phenylindole (DAPI) – 0,5 μg ml^-1^ (32670, Sigma-Aldrich; St. Louis, MO, United States); monodansylcadaverine (MDC) – 30 μg ml^-1^ (30432, Sigma-Aldrich) and acridine orange (AO) – 10 μg ml^-1^ (A6014, Sigma-Aldrich). To investigate an occurrence of apoptosis, In Situ Cell Death Detection Kit TMR red (12156792910, Roche, Basel, Switzerland) was used according to the manufacturer’s instructions. Preparations of cell samples for microscopy and the TUNEL assay were performed as described earlier (Lytvyn et al., 2010). Five fields of view (magnification 20X) containing more than 200 cells were examined in each microscopic analysis.

### CLSM

For investigation of autophagosome staining and cytoplasm acidification, plant samples were examined using a Zeiss LSM 510 META laser scanning confocal microscope equipped with lasers for 405 and 488 nm excitations, for MDC and AO, respectively. Images were collected using a 20x lens (EC Plan-Neofluar 20x/0,5, Zeiss) in multi-track mode with line switching and averaging of eights readings. Excitation of AO was performed in the different track equipped with 488 nm lasers and fluorescence for green and red emission spectra of AO was collected with a BP505–570 and LP650 band-pass filters, respectively. MDC was exited at 405 nm in the separate track and fluorescence was collected with a LP420 filter. Power of lasers was adjusted to 20% for excitation of 488 nm and to 15% for 405 nm, respectively. Estimations of 405/488 nm ratios of AO fluorescence were performed using Carl Zeiss Laser Scanning Microscope LSM510 Release Version 4.0 SP2. All experiments were performed in three replications. The results were presented as mean SE; statistical analysis was done by Student t-test and P values less than 0.05 were considered as statistically significant.

### Measurements of cellular proteins and glucose level

To determine changes in the level of total cellular proteins, 1-ml aliquots of cell suspension were pelleted at 500 g for 2 min, washed once with PBS, re-suspended in 1-ml of CelLytic^™^ P Cell Lysis Reagent (C2360, Sigma) and homogenised in liquid nitrogen followed by centrifugation at 13000 g for 20 min. The protein concentration was then measured according to the Bradford assay (Bradford, 1976) using BSA as a reference protein.

Conditioned cultural media of the collected samples were subjected to colorimetric evaluation of glucose concentration using Glucose (GO) Assay Kit (GAGO20, Sigma-Aldrich) following the manufacturer’s guidelines.

### PAGE and Western blotting

Lysates for electrophoretic analysis were prepared in the same way as for protein concentration measurement. Lysis buffer contained Protease Inhibitor Cocktail (P9599, Sigma-Aldrich) as well as 100 mM NaF, 1 mM Na_3_VO_4_, 100 nM okadaic acid. Proteins were separated by SDS-PAGE under reducing conditions and electro-transferred to nitrocellulose membrane (RPN3032D, GE Healthcare, Mickleton, NJ, United States) using the Bio-Rad Criterion Blotter. Before immunostaining membranes were pre-treated with Pierce Antibody Extender NC solution (32110, Thermo Fisher Scientific Inc., Waltham, MA, United States) according to the manufacturer’s protocol and blocked for 2 hours with 5% non-fat dry milk. Membranes were probed with monoclonal anti-acetylated tubulin (1:5000; T6793, Sigma Aldrich) or polyclonal anti-Glu-tubulin (detyrosinated) (1:5000; OBT1660, AbD Serotec/Bio-Rad Technologies) antibodies at 4°C overnight. Incubation with appropriate HRP-conjugated secondary antibodies was followed by detection with SuperSignal West Pico Chemiluminescent Substrate (34077, Thermo Fisher Scientific Inc.).

### Co-immunoprecipitation and Experion analyses

Co-immunoprecipitation was performed using the Pierce^®^ Co-Immunoprecipitation (Co-IP) Kit (26149, Thermo Fisher Scientific Inc.) according to the manufacturer’s recommendations. As was previously mentioned, anti-acetylated tubulin and anti-Glu-tubulin antibodies were for antigen binding. Antibodies were directly immobilised to aldehyde-activated beaded agarose (10 μg of antibodies to 50 μl of coupling resin per sample). Samples of cellular pellets harvested after 4, 11 and 14 days of cultivation (time periods that corresponded to exponential growth, autophagy and apoptosis-like stages) were used for bait:prey complex analysis. Pellets were lysed in supported IP-Lysis/Wash Buffer (3-ml buffer for approximately 300 mg of cells containing protease and phosphatase inhibitors) as mentioned above. Subsequently, the pellets were homogenised in liquid nitrogen and lysates were clarified by centrifugation at 13000 g for 10 min. 1 ml of lysate from each time point was incubated with 50 μl of prepared antibody-coupled resins at 4°C overnight. After specific incubation and washing procedures were performed, bound proteins were eluted in a total volume of 60 μl. Samples were pre-adsorbed to the control resin before co-immunoprecipitation to avoid non-specific protein binding to the coupling resin. Incubation of samples with coupled non-relevant antibody was performed as a negative control. Then, immunoprecipitates were analysed using the Experion automated electrophoresis system (Bio-Rad).

## ABBREVIATIONS

MT: microtubules
TUNEL: terminal deoxynucleotidyl transferase dUTP nick end labeling
PI: propidium iodide
DAPI: 4,6-diamidino-2-phenylindole
MDC: Monodansylcadaverine
AO: acridine orange (AO)

## COMPETING INTERESTS

The authors declare no competing or financial interests

## AUTHOR CONTRIBUTIONS

Dmytro I. Lytvyn – design and performing of experiments, data analysis and manuscript writing; Alla I. Yemets – performing of experiments and data analysis, Yaroslav B. Blume – data analysis and manuscript writing

## REFERENCES

Amos, L.A., Löwe, J. (1999). How Taxol^®^ stabilises microtubule structure. Chemistry and Biology. 6, 65–69.

Asada, T. and Shibaoka, H. (1994). Isolation of polypeptides with microtubule-translocating activity from phragmoplasts of tobacco BY-2 cells. Journal of cell science. 107, 2249–2257.

Avin-Wittenberg, T. Honig, A. and Galili G. (2012). Variations on a theme: plant autophagy in comparison to yeast and mammals. Protoplasma. 249, 285–299.

Baena-González, E., Rolland, F., Thevelein, J. M. and Sheen, J. (2007). A central integrator of transcription networks in plant stress and energy signalling. Nature. 448, 938–942.

Bass, B. P. Tanner, E. A, Mateos, D. Martín, S, Blute, T. Kinser, D. Dolph, P. J. and Mccall, K. (2010). Cell-autonomous requirement for DNaseII in non-apoptotic cell death. Cell Death Differ. 16, 1362–1371.

Bassham, D. C, Laporte, M., Marty, F., Moriyasu, Y., Ohsumi, Y., Olsen, L. J. and Yoshimoto, K. (2006). Autophagy in Development and Stress Responses of Plants. Autophagy. 2, 2–11.

Biederbick, A. and Kern, H. F. E. H. (1995). Monodansylcadaverine (MDC) is a specific in vivo marker for autophagic vacuoles. Eur J Cell Biol. 66, 3–14.

Boyault, C., Zhang, Y., Fritah, S., Caron, C., Gilquin, B., Kwon, S.H., Garrido, C., Yao, T. P., Vourc'h, C., Matthias, P. and Khochbin S. (2007). HDAC6 controls major cell response pathways to cytotoxic accumulation of protein aggregates. Genes Dev. 21, 2172–2181.

Boyle, K. M., Irwin, J. P., Humes, B. R. and Runge, S. W. (1997). Apoptosis in C3H-10T1/2 cells: roles of intracellular pH, protein kinase C, and the Na+/H+ antiporter. Journal of cellular biochemistry. 67, 231–40.

Bradford, M. M. (1976). A rapid and sensitive method for the quantitation of microgram quantities of protein utilizing the principle of protein-dye binding. Analytical biochemistry. 72, 248–54.

Cacas, J.-L. and Diamond. M. (2009). Is the autophagy machinery an executioner of programmed cell death in plants? Trends in plant science. 14, 299–300.

Cambray-Deakin, M. A. and Burgoyne, R. D. (1987). Acetylated and detyrosinated alpha-tubulins are co-localized in stable microtubules in rat meningeal fibroblasts. Cell motility and the cytoskeleton. 8, 284–291.

Chen. M. H., Liu, L. F., Chen, Y. R., Wu, H. K. and Yu, S. M. (1994). Expression of alpha-amylases, carbohydrate metabolism, and autophagy in cultured rice cells is coordinately regulated by sugar nutrient. The Plant journal: for cell and molecular biology. 6, 625–636.

Contento, A. L., Kim, S. and Bassham, D. C. (2004). Transcriptome Profiling of the Response of Arabidopsis Suspension Culture Cells to Suc Starvation. Plant Physiology. 135, 2330–2347.

Dunn, S., Morrison, E. E., Liverpool, T. B., Molina-París, C., Cross, R. A., Alonso, M. C. and Peckham, M. (2008). Differential trafficking of Kif5c on tyrosinated and detyrosinated microtubules in live cells. Journal of cell science. 121, 1085–1095.

Geeraert, C., Ratier, A., Pfisterer, S. G., Perdiz, D., Cantaloube, I., Rouault, A., Pattingre, S., Proikas-Cezanne, T., Codogno, P. and Poüs, C. (2010). Starvation-induced hyperacetylation of tubulin is required for the stimulation of autophagy by nutrient deprivation. The Journal of biological chemistry. 285, 24184–24194.

Giannoutsou, E., Galatis, B., Zachariadis, M. and Apostolakos, P. (2012). Formation of an endoplasmic reticulum ring associated with acetylated microtubules in the angiosperm preprophase band. Cytoskeleton. 69, 252–265.

Gottlieb, R. A., Giesing, H. A., Zhu, J. Y., Engler, R. L. and Babior, B. M. (1995). Cell acidification in apoptosis: granulocyte colony-stimulating factor delays programmed cell death in neutrophils by up-regulating the vacuolar H(+)-ATPase. Proceedings of the National Academy of Sciences of the United States of America. 92, 5965–5968.

Gundersen, G. G. and Bulinski, J. C. (1988). Selective stabilization of microtubules oriented toward the direction of cell migration. Proceedings of the National Academy of Sciences of the United States of America. 85, 5946–5950.

Halford, N. G., Hey, S., Jhurreea, D., Laurie, S., McKibbin, R.S., Paul, M. and Zhang, Y. (2003). Metabolic signalling and carbon partitioning: role of Snf1-related (SnRK1) protein kinase. Journal of Experimental Botany. 54, 467–475.

Hamada, T. (2007). Microtubule-associated proteins in higher plants. Journal of plant research. 120, 79–98.

Hammond, J., Cai, D. and Verhey, K. (2009). Tubulin modifications and their cellular functions. Curr Opin Cell Biol. 20, 71–76.

Hammond, J. W., Huang, C., Kaech, S., Jacobson, C., Banker, G. and Verhey, K. J. (2010). Posttranslational Modifications of Tubulin and the Polarized Transport of Kinesin-1 in Neurons. Molecular Biology of the Cell. 21, 572–583.

Han, S., Yu, B., Wang, Y. and Liu, Y. (2011). Role of plant autophagy in stress response. Protein & Cell. 2, 784–91.

Honig, A., Avin-Wittenberg, T., Ufaz, S. and Galili, G. (2012). A new type of compartment, defined by plant-specific Atg8-interacting proteins, is induced upon exposure of Arabidopsis plants to carbon starvation. The Plant Cell. 24, 288–303.

Huang, R. F. and Lloyd, C. W. (1999). Gibberellic acid stabilises microtubules in maize suspension cells to cold and stimulates acetylation of alpha-tubulin. FEBS Letters. 443, 317–320.

Hussey, P. J. and Hawkins, T. J. (2001). Plant microtubule-associated proteins: the HEAT is off in temperature-sensitive mor1. Trends in Plant Science. 6, 389–392.

Inoue, Y., Suzuki, T., Hattori, M., Yoshimoto, K., Ohsumi, Y. and Moriyasu, Y. (2006). AtATG genes, homologs of yeast autophagy genes, are involved in constitutive autophagy in Arabidopsis root tip cells. Plant & Cell Physiology. 47, 1641–1652.

Janke, C., Rogowski, K., Wloga, D., Regnard, C., Kajava, A. V., Strub, J. M., Temurak, N., van Dijk, J., Boucher, D., van Dorsselaer, A., Suryavanshi, S., Gaertig, J. and Eddé, B. (2005). Tubulin polyglutamylase enzymes are members of the TTL domain protein family. Science. 308, 1758–1762.

Khawaja, S. and Gregg, G. G. (1988). Enhanced Stability of Microtubules Enriched in Detyrosinated Tubulin Is Not a Direct Function of Detyrosination Level Quantxfication of Microtubules. The Journal of Cell Biology. 106, 141–149.

Kim, J., Kundu, M., Viollet, B. and Guan, K.-L. (2011). AMPK and mTOR regulate autophagy through direct phosphorylation of Ulk1. Nature Cell Biology. 13, 132–141.

Konishi, Y. and Setou, M. (2009). Tubulin tyrosination navigates the kinesin-1 motor domain to axons. Nature neuroscience 12, 559–567.

Li, J. and Eastman, A. (1995). Apoptosis in an interleukin-2-dependent cytotoxic T lymphocyte cell line is associated with intracellular acidification. Role of the Na(+)/H(+)-antiport. The Journal of Biological Chemistry. 270, 3203–3211.

Liu, Y. and Bassham, D. C. (2010). TOR is a negative regulator of autophagy in Arabidopsis thaliana. PloS One. 5. e11883.

Liu, Y. and Bassham, D. C. (2012). Autophagy: pathways for self-eating in plant cells. Annual Review of Plant Biology. 63, 215–237.

Liu, Y., Schiff, M., Czymmek, K., Tallóczy, Z., Levine, B. and Dinesh-Kumar, S. P. (2005). Autophagy regulates programmed cell death during the plant innate immune response. Cell. 121, 567–577.

Liu, Y., Xiong, Y. and Bassham, D. C. (2009). Autophagy is required for tolerance of drought and salt stress in plants. Autophagy. 5, 954–963.

Love, A. J., Milner, J. J. and Sadanandom, A. (2008). Timing is everything: regulatory overlap in plant cell death. Trends in Plant Science. 13, 589–595.

Lytvyn, D. I., Yemets, A. I., Blume, Y. B. (2010). UV-B overexposure induces programmed cell death in a BY-2 tobacco cell line. Environmental and Experimental Botany. 68, 51–57.

Mackeh, R., Perdiz, D., Lorin, S., Codogno, P. and Poüs, C. (2013). Autophagy and microtubules - new story, old players. Journal of Cell Science. 126, 1071–1080.

Mann, S. S., Hammarbacks, J. A., Gray, B. and Carolina, N. (1994). Molecular characterization of light chain 3. A microtubule binding subunit of MAP1A and MAP1B. The Journal of Biological Chemistry. 269, 11492–11497.

Monastyrska, I., Rieter, E., Klionsky, D. J. and Reggiori F. (2010). Multiple roles of the cytoskeleton in autophagy. Biol Rev Camb Philos Soc. 84, 431–448.

Nagata, T., Nemoto, Y. and Hasezawas, S. (1992). Tobacco BY-2 cell line as the ‘HeLa’ cell in the cell biology of higher plants. Int. Rev. Cytol. 132, 1–30.

Nakagawa, U., Kamemura, K. and Imamura, A. (2013a). Regulated Changes in the Acetylation of α-Tubulin on Lys40 during Growth and Organ Development in Fast Plants Brassica rapa L. Bioscience, Biotechnology, and Biochemistry. 77, 2228–2233.

Nakagawa, U., Suzuki, D., Ishikawa, M., Sato, H., Kamemura, K. and Imamura, A. (2013b). Acetylation of α-Tubulin on Lys40 Is a Widespread Post-Translational Modification in Angiosperms. Bioscience, Biotechnology, and Biochemistry. 77, 1602–1605.

Nezis, I. P., Shravage, B. V., Sagona, A. P., Johansen, T., Baehrecke, E. H., Stenmark, H. (2010). Autophagy as a trigger for cell death: Autophagic degradation of inhibitor of apoptosis dBruce controls DNA fragmentation during late oogenesis in Drosophila. Autophagy. 6, 1214–1215.

Nick, P. (2013). Microtubules, signalling and abiotic stress. The Plant Journal: for Cell and Molecular Biology. 75, 309–323.

Nishihama, R., Soyano, T., Ishikawa, M., Araki, S., Tanaka, H., Asada, T., Irie, K., Ito, M., Terada, M., Banno, H., Yamazaki, Y. and Machida, Y. (2002). Expansion of the cell plate in plant cytokinesis requires a kinesin-like protein/MAPKKK complex. Cell. 109, 87–99.

Patel, S., Caplan, J. and Dinesh-Kumar, S. P. (2006). Autophagy in the control of programmed cell death. Current Opinion in Plant Biology. 9, 391–396.

Perez, M., Santa-Maria, I., Gomez de Barreda, E., Zhu, X., Cuadros, R., Cabrero, J. R., Sanchez-Madrid, F., Dawson, H. N., Vitek, M. P., Perry, G., Smith, M. A. and Avila, J. (2009). Tau--an inhibitor of deacetylase HDAC6 function. Journal of Neurochemistry. 109, 1756–1766.

Reddy, A. S. N. and Day, I. S. (2001). Kinesins in the Arabidopsis genome: A comparative analysis among eukaryotes. BMC Genomics. 2, 2.

Reed, N. A., Cai, D., Blasius, T. L., Jih, G. T., Meyhofer, E., Gaertig, J. and Verhey, K. J. (2006). Microtubule acetylation promotes kinesin-1 binding and transport. Current Biology: CB. 16, 2166–2172.

Reumann, S., Voitsekhovskaja, O. and Lillo, C. (2010). From signal transduction to autophagy of plant cell organelles: lessons from yeast and mammals and plant-specific features. Protoplasma. 247, 233–256.

Rodionov, V. I., Lim, S. S., Gelfand, V. I., Borisy, G. G. (1994). Microtubule dynamics in fish melanophores. The Journal of Cell Biology. 126, 1455–1464.

Rolland, F., Baena-Gonzalez, E. and Sheen, J. (2006). Sugar sensing and signaling in plants: conserved and novel mechanisms. Annual Review of Plant Biology. 57, 675–709.

Rose, T. L., Bonneau, L., Der, C., Marty-Mazars, D. and Marty, F. (2006). Starvation-induced expression of autophagy-related genes in Arabidopsis. Biology of the Cell. 98, 53–67.

Sedbrook, J. C. (2004). MAPs in plant cells: delineating microtubule growth dynamics and organization. Current Opinion in Plant Biology. 7, 632–640.

Sheremet, Y. A., Yemets, A. I. and Blume, Y. B. (2012). Inhibitors of tyrosine kinases and phosphatases as a tool for the investigation of microtubule role in plant cold response. Cytology and Genetics. 46, 1–8.

Shimizu, S., Konishi, A., Nishida, Y., Mizuta, T., Nishina, H., Yamamoto, A. and Tsujimoto, Y. (2010). Involvement of JNK in the regulation of autophagic cell death. Oncogene. 29, 2070–2082.

Smertenko, A., Blume, Y., Viklicky, V., Opatrny, Z. and Draber, P. (1997). Post-translational modifications and multiple tubulin isoforms. Planta. 42, 349–358.

Smertenko, A. and Franklin-Tong, V. E. (2011). Organisation and regulation of the cytoskeleton in plant programmed cell death. Cell Death and Differentiation. 18, 1263–1270.

Telewski, F. W. (2006). A unified hypothesis of mechanoperception in plants. American Journal of Botany. 93, 1466–1476.

Tran, H. T., Nimick, M., Uhrig, R. G., Templeton, G., Morrice, N., Gourlay, R., DeLong, A., Moorhead, G. B. G. (2012). Arabidopsis thaliana histone deacetylase 14 (HDA14) is an α-tubulin deacetylase that associates with PP2A and enriches in the microtubule fraction with the putative histone acetyltransferase ELP3. The Plant Journal: for Cell and Molecular Biology. 71, 263–272.

Tsujimoto, Y. and Shimizu, S. (2005). Another way to die: autophagic programmed cell death. Cell Death and Differentiation. 12, 1528–34.

Verhey, K. J. and Gaertig, J. (2007). The Tubulin Code. Cell Cycle. 6, 2152–2160.

Walter, W. J., Beránek, V, Fischermeier, E. and Diez S. (2012). Tubulin acetylation alone does not affect kinesin-1 velocity and run length in vitro. PloS One. 7, e42218.

Wang, C., Li, J. and Yuan, M. (2007). Salt tolerance requires cortical microtubule reorganization in Arabidopsis. Plant & Cell Physiology. 48, 1534–1547.

Wang, W., Vignani, R., Scali, M., Sensi, E. and Cresti, M. (2004). Post-translational modifications of alpha-tubulin in Zea mays L are highly tissue specific. Planta. 218, 460–465.

Wang, C., Zhang, L.-J. and Huang, R.-D. (2011). Cytoskeleton and plant salt stress tolerance. Plant Signaling & Behavior. 6, 29–31.

Wasteneys, G. O. (2004). Progress in understanding the role of microtubules in plant cells. Current Opinion in Plant Biology. 7, 651–660.

Wingler, A., Masclaux-Daubresse, C. and Fischer, A. M. (2009). Sugars, senescence, and ageing in plants and heterotrophic organisms. Journal of Experimental Botany. 60, 1063–1066.

Xiao, W., Sheen, J. and Jang, J. C. (2000). The role of hexokinase in plant sugar signal transduction and growth and development. Plant Molecular Biology. 44, 451–461.

Xie, R., Nguyen, S., McKeehan, W. L. and Liu, L. (2010). Acetylated microtubules are required for fusion of autophagosomes with lysosomes. BMC Cell Biology. 11, 89.

Xie, R., Nguyen, S., Mckeehan, K., Wang, F., Mckeehan, W.L. and Liu, L. (2011). Microtubule-associated Protein 1S ( MAP1S ) Bridges Autophagic Components with Microtubules and Mitochondria to Affect Autophagosomal Biogenesis and Degradation. Journal of Biological Chemistry. 286, 10367–10377.

Xiong, Y., Contento, A. L. and Bassham, D. C. (2005). AtATG18a is required for the formation of autophagosomes during nutrient stress and senescence in Arabidopsis thaliana. The Plant journal: for Cell and Molecular Biology. 42, 535–546.

Xiong, Y. and Sheen, J. (2012). Rapamycin and glucose-target of rapamycin (TOR) protein signaling in plants. The Journal of Biological Chemistry. 287, 2836–2842.

Xiong, Y. and Sheen, J. (2014). TOR Signaling Networks in Plant Growth and Metabolism. Plant physiology, pp.113.229948.

Yano, K., Suzuki, T. and Moriyasu, Y. (2007). Constitutive Autophagy in Plant Root Cells. Autophagy. 3, 360–362.

Yemets, A. I., Krasylenko, Y. A., Lytvyn, D. I., Sheremet, Y. A., Blume, Y. B. (2011). Nitric oxide signalling via cytoskeleton in plants. Plant Science: an International Journal of Experimental Plant Biology. 181, 545–554.

Yoshimoto, K. (2012). Beginning to understand autophagy, an intracellular self-degradation system in plants. Plant & Cell Physiology. 53, 1355–1365.

Zelenin, A. V. (1966). Fluorescence Microscopy of Lysosomes and Related Structures in Living Cells. Nature. 212, 425–426.

